# The influence of selection, drift and immigration on the diversity of a tropical tree community

**DOI:** 10.1101/2022.06.27.497655

**Authors:** Jeronimo Cid, Ben Lambert, Armand M. Leroi

**Author notes:** joint senior authors & (Ben Lambert, Armand M. Leroi).

## Abstract

Ecology is rich in theories that aim to explain why natural communities have as many species as they do. Neutral theory, for example, supposes that a community’s diversity depends on the rate at which it gains species by immigration or speciation and loses them to ecological drift [1–5]. Classical niche theory, by contrast, supposes that diversity is regulated by the complexity of the environment: how many dimensions of resources it has and how finely species can subdivide them [6–10]. These theories are about levels of diversity at equilibrium. But non-equilibrium theory supposes that communities are perpetually buffeted by environmental change so that communities rarely contain all the individuals and species they might [11, 12]. When that happens, some species may profit from their immediate circumstances, but their gains are short lived as the environment changes again, favouring others. Such theories are often seen as competing visions of nature (e.g., [1, 2, 13–20]), but they can also be viewed as collectively describing a set of forces, any of which may be at work at a given time and place (cf. [21]). The relative importance of these forces in shaping the evolution of a community’s diversity can be captured by a small set of parameters: the community’s effective size, *N_e_*, the rate at which it gains new species, *μ*, and the magnitude and form of species-specific selection coefficients, *s* [22]. Here we present a way of estimating these parameters using time series data and apply it to the famous Barro Colorado Island Neotropical forest dataset. We show that, for the last thirty years, this community has been dominated by directional selection. We then simulate the evolution of this community in order to disentangle how these forces have shaped the species diversity that we see today. We show that, while species richness can be maintained by a neutral force, immigration, species evenness cannot and argue that it is likely maintained by temporally varying selection driven by environmental change [23–25].

---

Barro Colorado Island, Panama, contains a tropical tree community that is the source of a remarkable dataset. For over thirty years, beginning in 1982, every tree in a 50 ha study plot (“BCI”) — more than 400,000 of them distributed across 327 species — has been counted and identified at roughly five-year intervals [26]. Thus the dataset provides an excellent time-series of species abundances based on the identities of individual trees. Here, we use these time series data to estimate the relative importance of the forces that shaped species abundances at BCI between 1982 and 2015. To do this, we developed a general modelling framework in which the community observed in a given census year is determined by three processes: the reproduction and death of individuals present in the previous census year, the immigration of individuals belonging to new species, and ecological drift. This framework is the basis for three models, each of which assumes a different set of forces. We infer the relative importance of these forces by asking which of them best describes the data.

Our modelling framework assumes that, if *n_i_* is the number of individuals of the *i*th species, then, in the absence of selection and immigration, our best point estimate of the frequency of that species at time *t* + 1 is its frequency at time *t*. Ecological drift can, however, result in changes away from these expectations, and a multinomial distribution represents this process:

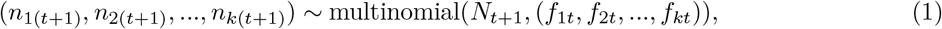

where *n_i_*(_*t*+1_) is the count of species *i* at time *t* +1, *k* is the count of species present at time *t*, *f_it_* = *n_it_*/*N_t_* is the frequency of species *i* at time *t*, and we take the total community size in each period 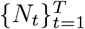 as given by the census data. Note that since the number of species which have appeared until a time *t* varies over time, *k* can change from year to year. See §0.3 for further details on modelling.

In our models, we allow selection to bias the frequencies of species in a child generation away from their parental values. We also introduce another term to account for the diluting effects of immigration of new species:

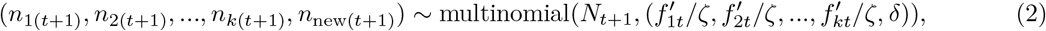

where *n*_new(*t*+1)_ is the number of individuals belonging to hitherto unobserved species which appear in period *t* + 1, and *ζ* is a normalising factor. Selection and immigration are embedded by their influence on the frequency of a species:

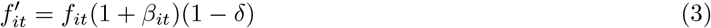

where *f_it_* = *n_it_*/*N_t_* is the frequency of species *i* at time *t*, and −1 ≤ *β_it_* ≤ 1 represents the strength and form of selection on each species; *δ* ∈ [0, 1] is the probability that an individual of a new species arrives in the community. This general framework resembles recent population genetic models that also use time series data [27–32], but differs from them in two ways. First, while population genetic models are built for two variants (biallelic SNPs), ours is built for many; second, our model does not take account of uncertainty in the estimation of species abundances or frequencies, but nor does it have to since, uniquely for BCI, these are known.

Our five models are variants of this framework that assume different forms of selection. Our first, and simplest, model assumes assumes that all *β_i_* are zero. Devoid of selection, it’s a pure neutral model in which the community’s abundance dynamics depend only on ecological drift and the immigration of new species. Our second model supposes selection is a function of frequency: *β_it_* = *ηf_it_*. This model speaks to niche-based theories because the signature of niches, conspecific negative density dependent growth (CNDD), manifests in communities as negative frequency dependent selection [21]. Our third model allows each species to have a separate, constant, *β_i_* value that is independent of frequency. This model speaks to non-equilibrium dynamics, because it estimates variation in competitive ability among species which, if present, implies a community that is not at a selective equilibrium but is, instead, in the throes of some disturbance or recovering from one. Our fourth and fifth models allow frequency independent selection with a selection strength, *β_it_*, which can vary over time: model 4 allows it to vary by census year; model 5 allows different selection strengths between the first and second half of observed census years (see §0.4 for more details).

Since our modelling framework assumes that all censused individuals are capable of reproduction, we filtered the BCI data to include only living adults. We excluded species in which we could not identify adults, and doing so left us with 169,747 individuals distributed across 258 nominal species. In an average census year there were *N* = 84, 384 ± 7039 (mean and 95%CI) adults, but their number declined slightly over the thirty three years of observation (Figure S1). Between one census and the next, an average of 11, 425 ± 2978 new adults were recruited and 13, 652 ± 2160 adults died, thus there was substantial turnover which, in turn, gave rise to the changes in species frequencies that we modelled. Model comparison revealed that the frequency independent model was a better fit to the data than any of the other models (see §0.7). Thus the abundance dynamics of the community are clearly not regulated by just neutral forces, and negative frequency dependent selection on adult trees, if present at all, must be much weaker than positive selection. This result does not exclude the possibility that negative frequency dependent selection acts on juveniles, nor does it allow that the strength of negative frequency dependent selection might vary among species [33].

Having established that BCI is dominated by frequency independent selection, we now focus on just that model. The selection parameters, *β_i_*, can be converted into absolute fitnesses, *W_i_* from which relative fitnesses, *w_i_*, and selection coefficients, *s_i_*, can be calculated (see §0.11). Selection coefficients ranged from 0.24 to −0.90. Of our 258 species, 57 — 22% — have selection coefficients whose 95% credible intervals do not embrace zero (Figure 1A). Since we expect to find 13 species (258 · 0.05) with such credibility intervals by chance alone, we conclude that some species are under positive or negative selection; indeed, these classes show the expected changes in frequency over time (Figure 1B). We considered the possibility that the observed variance in fitness among species was due to chance differences in their demographic composition rather than their intrinsic adaptedness, but further analysis allows us to reject this hypothesis (see §0.12).

**Figure 1:**
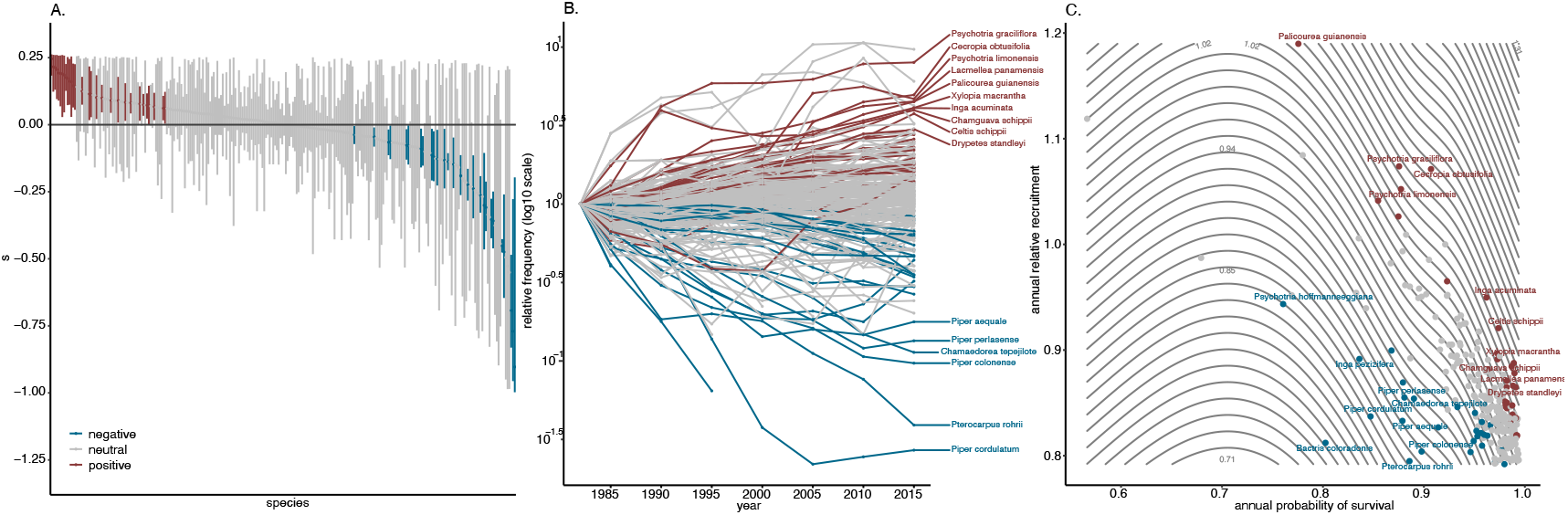
**A**. Frequency independent selection coefficients, *s_i_*, for 258 model species. *s_i_* are estimated from relative fitnesses which, in turn, are absolute fitnesses, *W_i_* relative to the median absolute fitness of all species (see §0.11). Species with a lower (2.5%) credible interval (CI) exceeding zero are marked in red and are deemed under positive selection; those whose upper (97.5%) CI was less than zero are marked in blue and are deemed under negative selection; the remainder, whose dynamics we cannot distinguish from neutrality, are marked in grey. **B**. The estimated selection coefficients predict the observed relative changes in frequency; ten examples of species estimated as undergoing positive selection are labelled along with seven undergoing negative selection. **C**. The relationship between two fitness components, annual relative recruitment rate, *RR_i_*, and the annual probability of survival, *SR_i_*, and relative fitness, *w_i_*, as estimated by a generalised additive model through the *gam* function within the mgcv R package [34]. Contour lines in grey show the estimated fitness surface.

Variation in relative fitness among species could be due to either variation in their average survivorship, variation in the average number of adult progeny that surviving individuals have, or both. So we next asked which of these two fitness components mattered most. The BCI records are based on individual trees, so we know when each tree was seen first and when it was seen last. We could therefore use these data as the basis of a new model in which we estimated the annual survival rates and recruitment rates for each species (see §0.5 for model details). We estimated the annual survival rate for each species, *SR_i_*, by assuming that trees died in the census interval after they were last seen. The species with the highest *SR* was *Hura crepitans* (0.994; 95% credibility interval: 0.991–0.996) and the lowest *Clidemia octona* (0.565; 0.439–0.673) implying that these species respectively lost about 0.6% and 44% of their adults each year due to mortality. We also estimated the annual recruitment rate of each species relative to the number of individuals present, *RR_i_*, by assuming that each tree started reproducing in the census interval after it first reached reproductive size. The species with the highest *RR* was *Palicourea guianensis* (1.19; 1.131–1.211) and the lowest was *Astrocaryum standleyanum* (0.792; 0.79–0.795). This implies that, ignoring adult mortality, each year these species gained and lost about 20% of adults respectively relative to those present. We then used a linear model, with an interaction term, to estimate the relative contribution of these fitness components to relative fitness, *w_i_*, as estimated above. We found that all terms were highly statistically significant (*P* < 2.0 · 10^−16^), that together they explained 72% of the variance in relative fitness, and that *SR_i_* accounted for 51% of the explained variance, while *RR_i_* accounted for 28% and their interaction about 21%. Thus, differential mortality appears to be the primary selective force shaping species abundances. Figure 1C shows the relationship among these variables overlain by a fitness surface estimated using a generalised additive model that explained 96% of the variance. The most successful — fittest — species are located North-East of the *w* =1 contour line, and the least South-West of it. Interestingly, it appears that successful species are not all successful in the same way. Where *Palicourea guianensis* has a high recruitment rate, but a mediocre survivorship, the opposite is true for *Drypetes standleyi*. On the other hand, the wild coffee, *Psychotria hoffmannseggiana*, has both low *SR* and *RR* and accordingly low fitness.

We also used this model to estimate the annual rate at which new species arrived, relative to the number of individuals present, as *δ* = 1.19 · 10^−4^ (95% Credible Intervals: 5.60 · 10^−5^ – 2.16 · 10^−4^) which implies that the community, as represented by this data set of adults, gained a new species about every 5.50 years (95% CI: 2.32-22.00 years), or 6.00 (95% CI: 1.50-14.25) species over the entire series, which agrees well with the 6 new species actually observed. Our estimate of the rate of arrival of new species is about half that estimated by [26], either because we only counted adults where they also counted juveniles or else because we did not allow species to immigrate multiple times after extinction, where they did.

Having estimated the forces that influence species abundances at BCI, we next asked which of them most influenced diversity. Between 1982 and 2015, species diversity at BCI changed rapidly. Species counts, ^0^*D*, show that, despite the immigration of new species, the community lost about one species every two years (Figure 2A). At the same time the effective number of species, as estimated by Simpson’s index of diversity, ^2^*D*, increased at rate of one effective species every twenty two years. Thus species abundances became more even. These estimates are based the data set of 258 species of adults used in our selection analysis; the complete data set of 327 species, which included juveniles, shows a qualitatively similar pattern except that the rate of species loss was slower and the rate of increase of species evenness faster (Figure S7).

**Figure 2:**
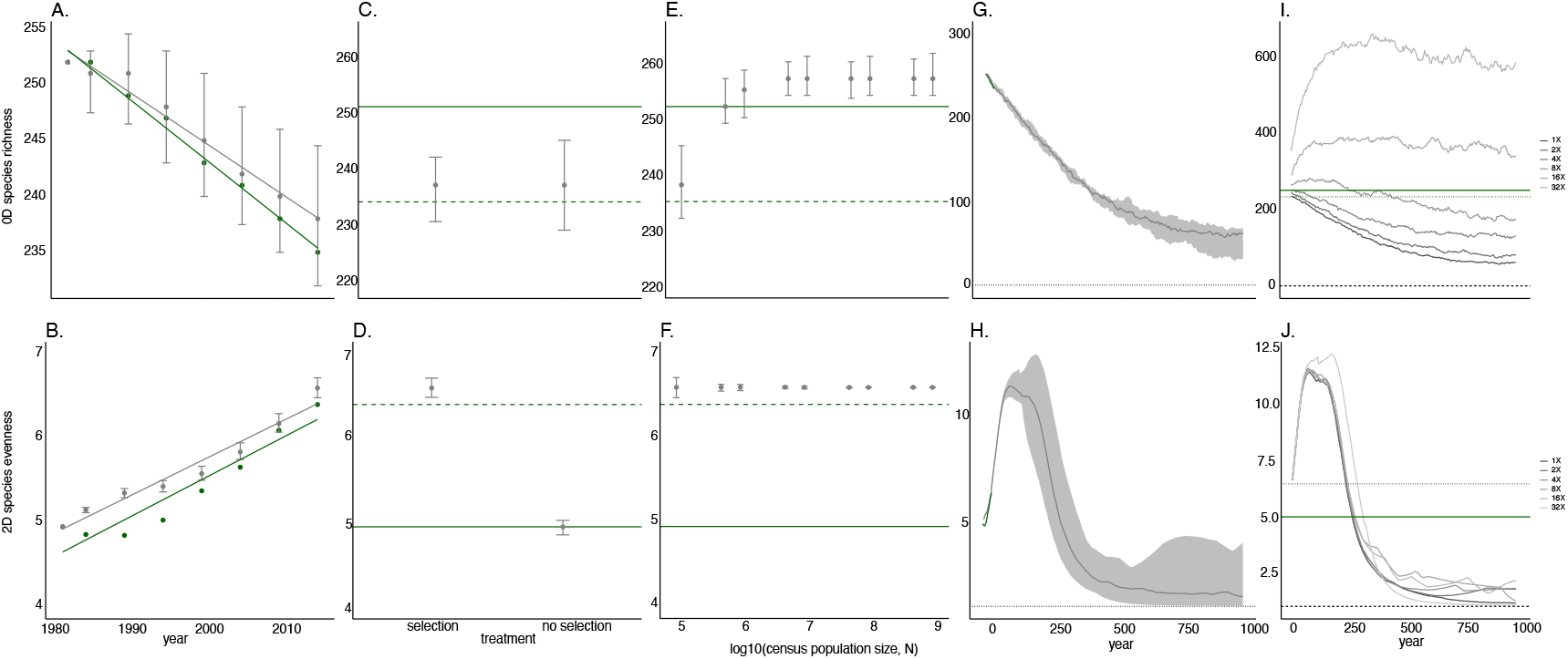
The regulation of diversity. **A**. Between 1982 and 2015 species richness declined at BCI at a rate of −0.537 ± 0.052 (estimate from linear regression model and 95% Confidence Interval) species per year (green). In our simulated communities, species richness declined at very similar rate of −0.453 ± 0.073 (grey). **B**. At the same time, species evenness increased at a rate of 0.049 ± 0.016 effective species per year (green), while in our simulated communities (grey), species evenness increased at a very similar rate of 0.045 ± 0.01. **C**–**F**. The effect of eliminating selection and drift on the diversity of simulated BCI communities 1982–2015; points show the median diversity of simulated communities at 2015 (± empirical 95% CIs), while the horizontal lines show the diversity present in the actual community in 1982 (solid green line) or 2015 (dashed green line). **C**. Removing selection has a no detectable effect on species richness (*P* = 0.316). **D**. Eliminating selection has a considerable effect *P* < 10^−16^) on species evenness so that the 1982 level of evenness is maintained in 2015. **E**. Removing drift by increasing the census community size, *N*, has a considerable effect on species richness up to about 4m trees, so that 1982 levels of species richness are maintained in 2015 (*P* < 10^−16^). **F**. Eliminating drift does not have a statistically significant effect (*P* = 0.423) on species evenness. All *P* values based on linear models. **G**–**J**. The long term evolution of simulated BCI communities projected for a millennium after 2015. These simulations assume the community size, selection coefficients and migration rates of new species that were estimated for BCI 1982-2015. Dotted horizontal lines mark a diversity of 1; solid lines are medians; shaded ribbons are 95% empirical CIs. **G**. The long-term evolution of species richness of simulated BCI communities. Species richness declines monotonically, continuing the observed trend from 1982–2015. **H**. The long-term evolution of species evenness in the same communities as panel G. After an initial increase species evenness declines steadily below the 1982 value. **I**. Simulated communities showing that increasing the immigration rate by a factor between four and eight can maintain species richness at the 1982 levels indefinitely. **J**. The same simulated communities in panel I showing that increasing the immigration rate cannot maintain species evenness, indeed, hastens its decline.

To disentangle the forces responsible for these changes in diversity we built a simulation of the BCI tree community based on our statistical model incorporating survival and recruitment (see §0.10). This simulation began with species abundances as they were in 1982 and projected forward until 2015 and assumed that the abundance of each species depended on its annual survival rate, *SR*, and relative recruitment rate, *RR*. It also assumed the observed community size at each census year, the community’s estimated immigration rate, and incorporated ecological drift via a multinomial sampling process among census years.

In order to use our simulation to test causal hypotheses, we need to show that it captures the dynamics of the real community. Figure S8 compares the simulated and actual frequency dynamics of the nine most abundant species. For most species the simulated frequencies track the observed frequencies. A few, however, have abundance trajectories that reversed direction dramatically at some point, and the simulation was less successful in tracking these. This is because our simulation assumed a single value for each fitness parameter for each species when in fact they may have varied over time in the wild. Such fluctuations in abundance, then, hint that the direction of selection may not have been constant at BCI (although models including fluctuating selection did not perform as well as the frequency independent model: see §0.7). Nevertheless, when we estimated the relative change in abundance of species between 1982 and 2015 for both the observed and simulated communities we found that they were highly correlated (*r*^2^ = 0.88) which shows that the simulation successfully recapitulated the overall frequency dynamics of the community over the course of 33 years (Figure S9). Since our simulation was based on frequency independent selection only, this is further evidence that frequency dependent selection at any age cannot be a major force shaping abundance dynamics. Most importantly, our simulation recapitulates the observed diversity dynamics BCI between 1982 and 2015 remarkably well (Figures 2A and B).

To test the influence of selection and drift on diversity we carried out several experiments in which we eliminated either selection or drift from our simulation. Selection was eliminated by setting the annual survival rate, *SR*, and the relative recruitment rate, *RR*, for all species to a constant, the average of all species. The effect of drift was reduced by increasing the observed community size by a factor of 1, 5, 10, 50, 100, 500, 1000, 5000 or 10000 so that the largest treatment had, in any census year, 8.36 · 10^8^ trees.

We hypothesized that selection, rather than drift, was the main cause of the decrease in species richness in BCI between 1982 and 2015. We thought that eliminating selection from our simulation would increase species richness since previously unfit species that might have gone extinct would no longer do so. Conversely, since the effect of drift on the frequency dynamics scales as as 1/*N_e_*, and since the adult census community size of BCI is already large (> 10^4^ individuals), we thought that reducing drift further should have very little effect. These predictions were wrong. Eliminating selection has essentially no effect on species richness in 2015 (Figure 2C), while reducing drift progressively increases species richness until it plateaus just above the 1982 level (Figure 2E). Thus, ecological drift, rather than selection, appears to have been the main cause of the observed decline at BCI.

We also hypothesized that selection, rather than drift, was the main cause of the increase in species evenness in BCI between 1982 and 2015. In contrast to our species richness results, these predictions were confirmed: increasing community size in our simulation has no detectable effect on species evenness, but eliminating selection keeps the community at its low 1982 level (Figures 2D, F). We conclude that, even though the change in the number of species observed at BCI was regulated by neutral forces, their change in relative abundance was regulated by selection.

In the absence of new migrant species, frequency independent selection and drift will cause a community to eventually become a monoculture. Even when immigration is present, they may cause a community to become an effective monoculture, that is, be so dominated by one species that the species evenness index, ^2^*D* approaches 1. To evaluate the long-term consequences of the forces working on BCI’s community dynamics, we ran the simulation forward using the selection coefficients and immigration rate that we estimated and a uniform community size of 83,648 individuals based on the harmonic community size 1982-2015. The survival rates and relative recruitment of new migrant species were drawn at random from a distribution based on the existing species (see §0.10).

This forward simulation showed the previously observed rise in species evenness (Figure 2B) continuing for about a century. A precipitous decline follows, however, so that within about 500 years the community contains only two effective species (^2^*D* = 1.93; LCI:1.2; UCI:3.31; 95%CI) (Figure 2H). The two most abundant species at this time — typically *Chamaguava schippi* and *Xylopa macrantha* — have a combined median frequency of 0.83; by contrast the two most abundant species in 1982, *Hybanthus prunifolius* and *Faramea occidentalis*, have a combined frequency of just 0.50. In our simulated community, *C. schippii* continues to increase in frequency until it dominates (Figure S10). Plainly it is a very fit species. Given the uncertainties of our fitness estimates we cannot, however, say that it is the fittest.

Thus the observed increase of ^2^*D* in the face of directional selection is merely a transient phenomenon due to the decrease in frequency of some of the most common species which just happen to be under negative selection. How much immigration is needed to counter the long-term diversity-depleting effects of selection and drift? To find out we ran simulations with increasingly high values of immigration and found that, to maintain species richness, ^0^*D*, at 1982 levels in the long term required between four and eight times the actual number of new species that arrive at BCI annually (Figure 2I). Thus species richness can be maintained indefinitely by modest increases in immigration. This is not true, however, of diversity in the sense of species evenness for, regardless how much we boosted immigration, ^2^*D* ultimately crashed (Figure 2J). Increasing immigration even hastened the rate at which species evenness declined, since it increased the probability of arrival of a new migrant species superior to any existing one.

The purpose of our forward simulation is not so much to offer a prediction of what BCI will look like in the future, say in the year 2500 CE, but rather to test the uniformitarian assumption that the forces currently present can account for the diversity that we see. The extraordinary diversity of tropical tree communities has usually been explained by either appealing to neutral forces or else niche-based selection [14, 35]. Our results, however, show that neither explanation is adequate, for the main force that has shaped species abundances at BCI over the last thirty or so years is directional selection. Our forward simulation shows, furthermore, that the observed levels of directional selection have the power to turn BCI into a community composed of one or two dominant species and an ever-changing repertoire of rarities — in just centuries. Neither frequency dependent selection nor immigration can counter this homogenizing force, for the first appears to be negligible, and the second has little effect on species evenness. Indeed, if the forces that we have estimated are typical of those acting on a patch of forest over the last few thousand years, we may wonder why BCI, or the Neotropical forest as whole, isn’t a near-monoculture already.

One answer is that the selection regime that we identified was preceded by others so that the diversity that we observe today is the result of a long history in which different species, with different ecological needs, have been favoured at different times. This is the “non-equilibrium” view of community structure championed by Connell [11] more than forty years ago. Indeed, previous studies of BCI and other communities, although not demonstrating selection, have shown that species abundances are driven, in part, by environmental stochasticity [23, 25, 36]. Our results hint that selection coefficients have not been constant over even thirty years — the generally higher mortality and reduced recruitment in the late 1990s–early 2000s suggests a particularly trying period for BCI’s trees (Figure S1). We examined this possibility explicitly by testing frequency independent models that allowed temporal variation in selection coefficients against those that did not, but found that they were poorly supported by the data (see §0.7). Given that we only have data on seven time points such tests, however, are probably not very powerful. We also note that, although BCI is assumed to be a model of a patch of Neotropical forest, it may in fact be rather unusual. Barro Colorado Island was formed only in 1913 by the flooding of the Panama canal, so the strong directional selection that we observed may be an echo of environmental changes [37] not found elsewhere.

There is another possibility: that it is not BCI that is unusual, but our times. Previous studies have documented the changes in species abundances that form the basis of our selection estimates and linked them to changes in weather patterns such as El Niño events [24, 38, 39]. Such changes are part of a more general increase in the abundances of drought-tolerant species in Neotropical forests [40, 41]. If climate change is indeed the selective agent, and if the rate of climate change remains unchecked, then our findings do not so much speak to its past but its future, and predict that BCI will be essentially a monoculture just a few centuries from now.

## Acknowledgments

We thank J. Rosindell, R. Chisholm, G. Bell and A. Burt for useful discussions.

## Supplementary Materials and Methods

### 0.1 Data and code availabilty

Our data, R, Stan and Julia code are available in a [GitHub repository](https://github.com/ben18785/bci_temporal_dynamics), which can be used to run the analysis from start to finish via GNU Make.

### 0.2 Data

The basis of our study is census data from the 50 ha. Barro Colorado Island plot [42, 43]. Our data contains counts in seven census years: 1982, 1985, 1990, 1995, 2000, 2005, 2010, 2015, that is, 4.7 years apart on average. We wanted a dataset that consisted only of living adults. To obtain living trees, we filtered the data for `status = A` for “alive”. To obtain adults we made use of a dataset which gave the minimum size at reproduction (`robin`) in terms of `breast height diameter` (`mm`) (`bhd`) for many species and only counted trees for which `bhd` ≥ `robin`. To simplify the modelling, we also excluded some trees that belonged to species that went extinct and then reappeared again. For example, *Psychotria hoffmannseggiana* was present in 1982, absent in 1985, present in 1990, absent in 1995, 2000 and 2005 and then present again in 2010 and 2015; but we ignored these cases and assumed that, once a species was extinct it was forever so. This left us with 169,747 individuals distributed over 258 species.

Since many tree species mature slowly and are long lived it may be supposed that the BCI study plot was — at least on the time scale of the study, a few decades — demographically inert. The total number of adults present didn’t change much: in 1982 there were 89,769; in 2015, 74,186, implying an annual loss of about 660 per year (Figure S1). In our dataset, 79,978 individuals were born after 1982 and 95,561 individuals died before 2015 (Figure S1).

**Figure S1:**
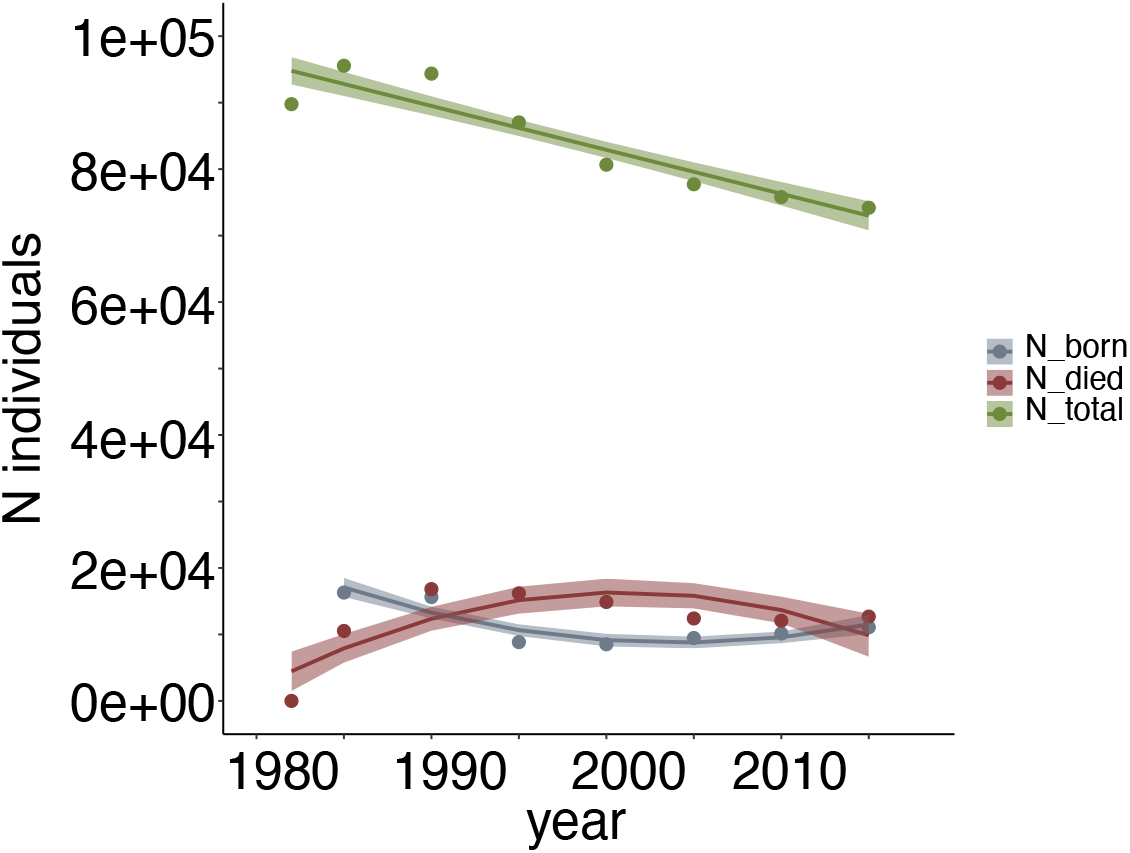
Total abundance, births and deaths of adults in the BCI 50 ha. study plot, 1982-2015. Total abundances; a linear model (fit ± 95%CI) shows a decline of −660.5 ± 261 adults per year (*P* = 0.0008). For annual births and deaths, quadratic models provided a superior fit suggesting that the change in the annual rate of births and deaths was not monotonic. Between 1995 and 2000, recruitment seems have to been particularly poor and deaths exceptionally high. The uncertainty ribbons indicate the standard error in the fits.

### 0.3 Modelling the frequencies of species in a zero-sum community over time

We modelled the dynamics of BCI’s trees in frequency space. There are two basic neutral models available: Wright-Fisher and Moran. The former assumes that, at each time point, new trees are sampled at random from parents, and that all trees die each time point and so all are replaced. The second assumes that, at each time point, new trees are sampled at random from parents, but that only one tree dies and is replaced. It’s clear that neither accurately describes the BCI situation where, between time points (census years), about 14% of trees are new. For this reason, we construct models based on the observed turnover of individuals.

#### Defining immigrants and residents

We need to distinguish where new individuals come from. Individuals observed for the first time in any census after 1982 are either *new residents* or *immigrants*. If an individual observed for the first time at *t* + 1 belongs to a species present in the plot at *t*, then we assume that it is the offspring of an individual in the plot: it’s a *new resident*. If a new individual belongs to a species never before seen in the plot, or that was previously observed but is extinct at *t*, then we assumed that it is an *immigrant*.

### 0.4 Models of temporally varying frequency independent selection

Eqs. (2)&(3) dictate the dynamics of variant frequencies within our models. Our fourth and fifth models allow selection strength, *β_it_* to vary in a frequency independent fashion, but they also allow *β_it_* to vary over time.

Model 4 allows *β_it_* to vary by census year, and we use the following random walk prior for this quantity:

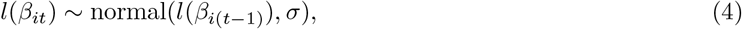

where *l*(*β*) is a transformation of *β*, so that it is on the unconstrained scale and is given by the inverse of: 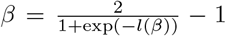. The use of such a random walk prior is to smooth the changes in the selection coefficients from one period to the next.

Model 5 allows two values of selection strength for each species: one before 1995 (*β*_*i*1_) and one after 1995 (*β*_*i*2_).

### 0.5 Birth and death model

Here, we describe how annual survivorship and recruitment were estimated at the species level. To estimate survivorship, we used a binomial sampling model:

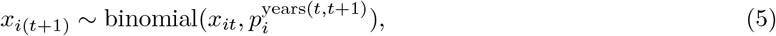

where *x*_*i*(*t*+1)_ is the count of those individuals of species *i* which remained in census year *t* + 1 after being recorded as present in the previous census year; 0 ≤ *p_i_* ≤ 1 is the annual probability of survival for that species; and years(*t,t* + 1) indicates the number of years between census years *t* and *t* +1.

To estimate recruitment, we used a model similar in spirit to the overall multinomial model:

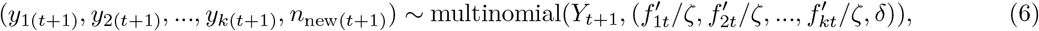

where *y*_*i*(*t*+1_) indicates the number of previously unobserved trees of species *i* which appear in census year *t* +1; *n*_new(*t*+1)_ is the number of new trees of species hitherto unseen in BCI which appear in the same year; *Y*_*t*+1_ is the total number of newly recorded trees that appear; 0 ≤ *δ* ≤ 1 is a probability that determines the rate of appearance of individuals of new species; and 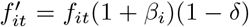, where, here, *f_it_* indicates the parental frequency of species *i* in census year *t*, and −1 ≤ *β_i_* ≤ 1 is a parameter representing how selection mediates recruitment; 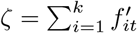 is a normalising factor.

#### Uneven census years

Eqs. (2)&(6) assume that the census years are evenly spaced. In our dataset, the census years were unevenly distributed, with five year gaps between all but one pair of consecutive censuses, where it was three, in 1982-1985. To account for this, in the models with selection, we replaced (1 + *β_i_*) → (1 + *β_i_*)^3/5^ and 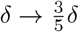 for the pair of years with this shorter gap.

**Table S1:** Summary of priors assumed in each analysis. Note, that the random walk prior for the corresponding model is given by eq.0.4.

| model | equation | priors |
| --- | --- | --- |
| neutral | eq. (2) | $\delta \sim U(0, 1)$ |
| frequency independent | eq. (2) | $\beta_i \sim U(-1, 1)$<br>$\delta \sim U(0, 1)$ |
| frequency dependent | eq. (2) | $\eta \sim U(-1, 1)$<br>$\delta \sim U(0, 1)$ |
| random walk | eqs. (2) & (4) | $\sigma \sim \text{normal}(0, 0.1)$<br>$l(\beta_{i0}) \sim \text{normal}(0, 0.5)$ |
| split selection strength | eq. (2) and §0.4 | $\beta_{it} \sim U(-1, 1), i \in \{1, 2\}$ |

### 0.6 Model priors

The models were estimated in a Bayesian framework. As such, the parameters were each given priors, which are provided in Table S1 for the non-birth-death models (i.e. not those models described in §0.5). The priors were chosen to be uninformative with the exception of the prior for *σ* in the random walk model (see §0.4), which was chosen to constrain the degree to which selection strength could vary over time.

For the birth-death model described in §0.5, we assumed the following bivariate hierarchical prior for each species *i* to account for the likely negative correlation between recruitment and mortality rates:

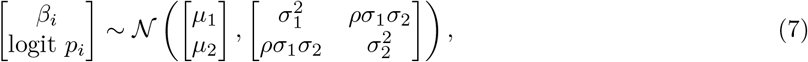

where logit *z* = log(*z/*(1 – *z*). The variables (*μ*_1_, *μ*_2_, *ρ*, *σ*_1_, *σ*_2_) were assumed to be common across all species, and for these, the following priors were specified:

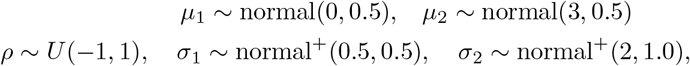

where normal^+^ indicates a normal distribution that is truncated at zero and has non-zero probability densities only for non-negative values. For this model, we performed prior predictive checks, which resulted in the uninformative joint distribution shown in Figure S2, where the marginal distributions had the following 2.5%-97.5% quantiles: −0.90 ≤ *β_i_* ≤ 0.91 and 0.19 ≤ *p_i_* ≤ 0.999.

### 0.7 Model comparison

To compare the predictive power of the different models covered by eq. (2), we performed explicit leave-one-out cross-validation. In this process, we fit the model to all census years bar one and determined the log-likelihood when predicting the held out year of data. This procedure was repeated across all five nonbirth-death models considered, and the results are shown in Table SS2. This indicates that the frequency independent model had the best predictive fit of all the models considered.

### 0.8 Model fitting

The models were estimated via Markov chain Monte Carlo (MCMC) using Stan’s NUTS algorithm using default settings [44]. In all cases, 4 Markov chains were used. For the models described by eq. (2) (including the models fit on a subset of the data when performing model comparison as described in §0.7) between 4000 and 16,000 iterations were used, and chains were thinned by factors ranging between 2-20 (see our [GitHub repository](https://github.com/ben18785/bci_temporal_dynamics) for details of individual model MCMC specifications). For the birth-death model described in §0.5, the MCMC was run for 200,000 iterations, and the chains were thinned by a factor of 100.

In all cases (including the hold-out fits described in §0.7) across all parameters, 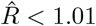, and the bulk- and tail-ESS values were above 400 diagnosing MCMC convergence [45].

**Figure S2:**
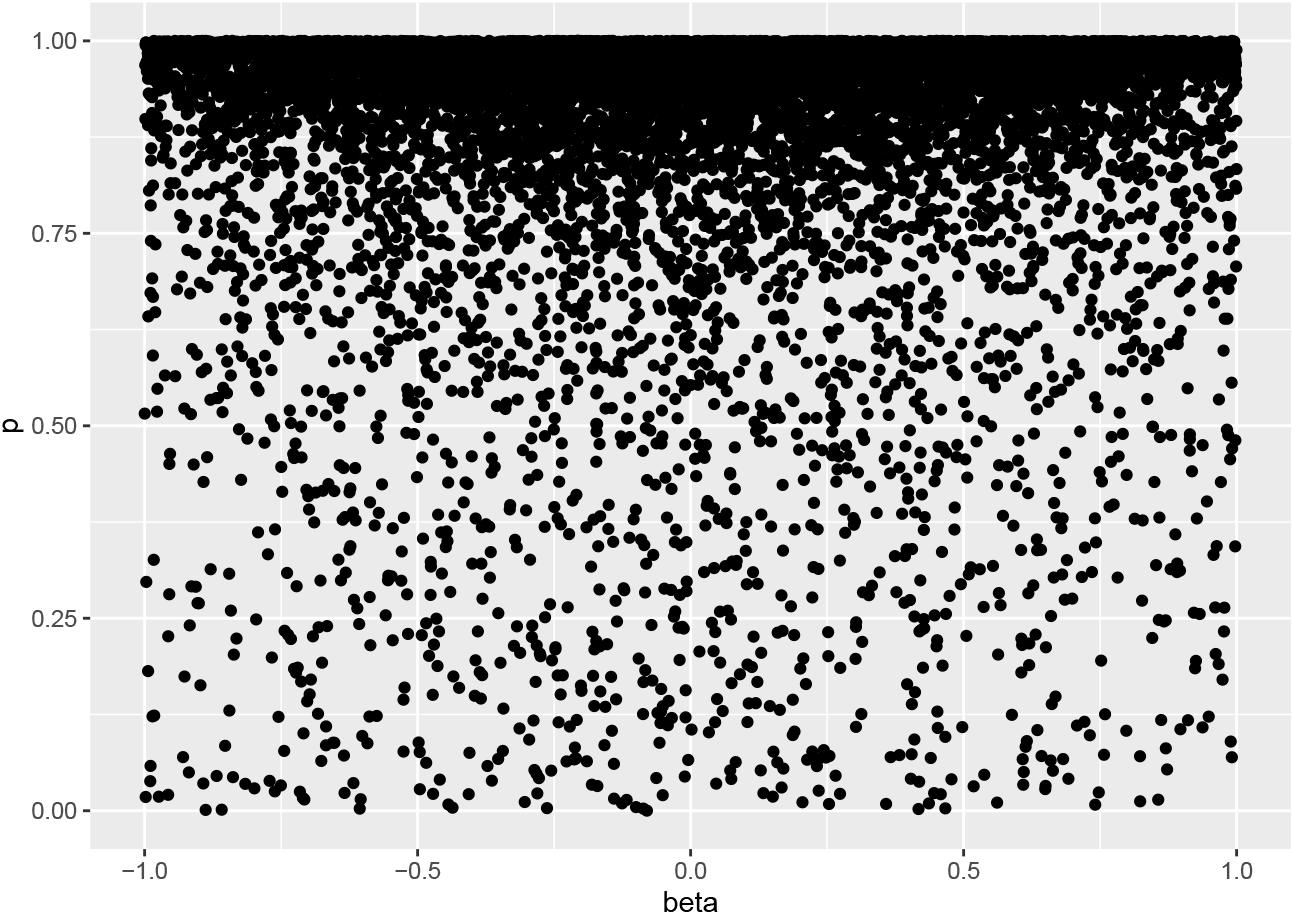
Prior predictive distribution for birth-death parameters. This was generated by drawing 10,000 values from the hierarchical prior distribution given by eq. (7).

**Table S2:**
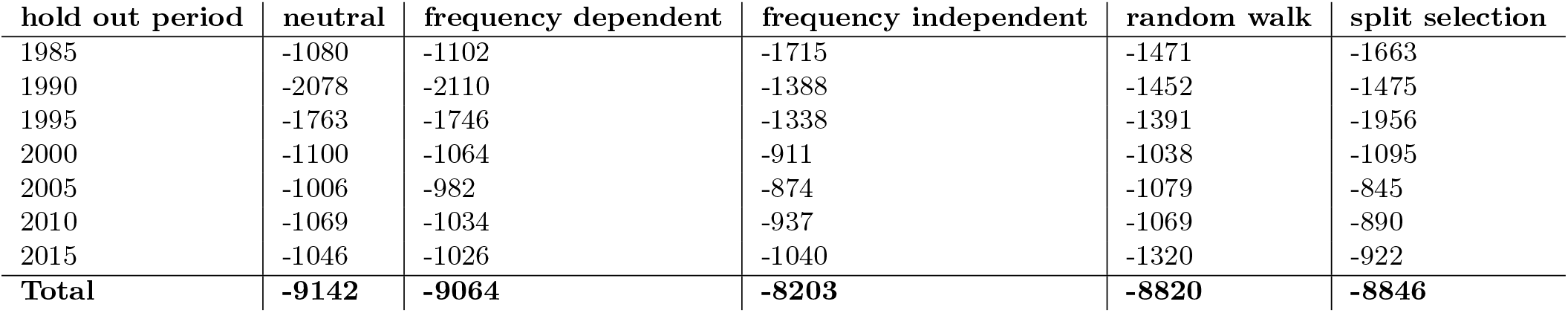
Model comparison: hold out predictive accuracies. Each row (apart from the bottom) indicates the predictive accuracy (in terms of the log-likelihood averaged over all posterior draws) for each model for the hold out period’s data. The bottom row shows the total of the mean log-likelihoods across all hold out periods.

| hold out period | neutral | frequency dependent | frequency independent | random walk | split selection |
| --- | --- | --- | --- | --- | --- |
| 1985 | -1080 | -1102 | -1715 | -1471 | -1663 |
| 1990 | -2078 | -2110 | -1388 | -1452 | -1475 |
| 1995 | -1763 | -1746 | -1338 | -1391 | -1956 |
| 2000 | -1100 | -1064 | -911 | -1038 | -1095 |
| 2005 | -1006 | -982 | -874 | -1079 | -845 |
| 2010 | -1069 | -1034 | -937 | -1069 | -890 |
| 2015 | -1046 | -1026 | -1040 | -1320 | -922 |
| **Total** | **-9142** | **-9064** | **-8203** | **-8820** | **-8846** |

### 0.9 Posterior predictive checks

In Figure S8, we show the actual and simulated frequency dynamics of the nine most common species when usin the frequency independent model. This indicates there the model was often able to account for variation in species frequency across all census years, but, inevitably, there were cases (since this model does not allow temporal fluctuations in selection) where the model was unable to recapitulate some of the more complex observed dynamics.

For the birth-death model, we performed a range of posterior predictive checks of the model’s fit. We compared the simulated probability of survival versus the actual for each species. First, we show this information when sorting these quantities according to the number of parents present in a given census year (Figure S3). This indicates that, as the number of parents increased, our model was better able to determine the survival rates. We also compared the simulated and actual survival rates for each census year (Figure S4). Across all census years, this indicated that the model produced reasonable predictions. We also compared the simulated and actual numbers of offspring of each species (Figure S5), which showed strong correspondence between these quantities. In addition, the results shown in Figures 2A & B indicate that simulated species richness and species evenness tracked the observed.

**Figure S3:**
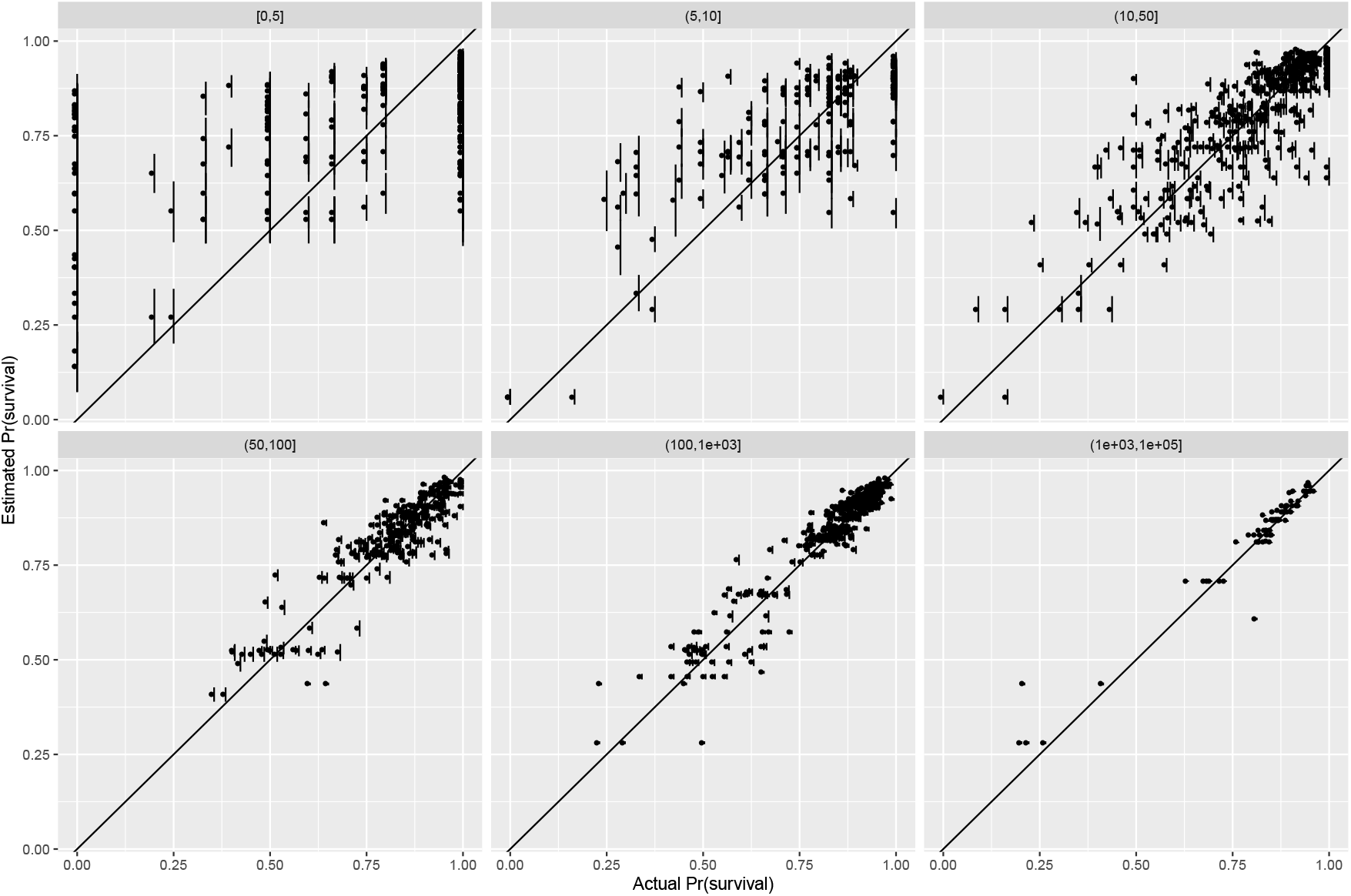
Posterior predictive distribution for the birth-death model: annual survival probabilities by number of parents. Each panel indicates actual versus predicted annual survival probabilities for a given range of numbers of parents present in the previous census year. The vertical point-range whiskers indicate 25-75% posterior quantiles; the vertical point positions indicate posterior medians.

### 0.10 Simulating BCI

To simulate the abundance of trees in BCI, we used the model described in §0.5 where the parameters were estimated by fitting the model to data from 1982-2015. To generate the number of migrant individuals, we used a Poisson distribution approximation:

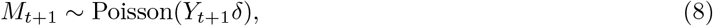

where, as in §0.5, *Y*_*t*+1_ is the total number of newly recorded trees that appear, and 0 ≤ *δ* ≤ 1 is an estimated probability that determines the rate of appearance of individuals of new species. All migrants were assigned to new species, but, like in the observed data, each new migrant was not necessarily a unique species. In the observed BCI data, there were 1.33 migrant individuals per census year, but this corresponded to only 0.825 species per census year. We used these estimates to allocate our migrants to new species by multinomial sampling with uniform probabilities. For a new species, its (*β_i_, p_i_*) parameters governing its reproductive and survival fitnesses were drawn from the hierarchical distribution described by eq. (7).

When simulating the model for the observed years 1982-2015, we supposed that the total number of trees and the number of newly arriving individuals in each census year from 1982-2015 were known. When extrapolating outside of this range (until the year 3000), the overall community size was set to the harmonic mean across observed census years (83,648 individuals). The interval between time-steps beyond 2015 was set to the average census interval for the series (4.125 years/census). The average offspring per tree per time-step was set to the average observed (0.13 offspring per parent), meaning the number of newly arriving individuals was 10,874 in each census year.

Depending on the experiment, we ran between 5-100 replicate simulations.

**Figure S4:**
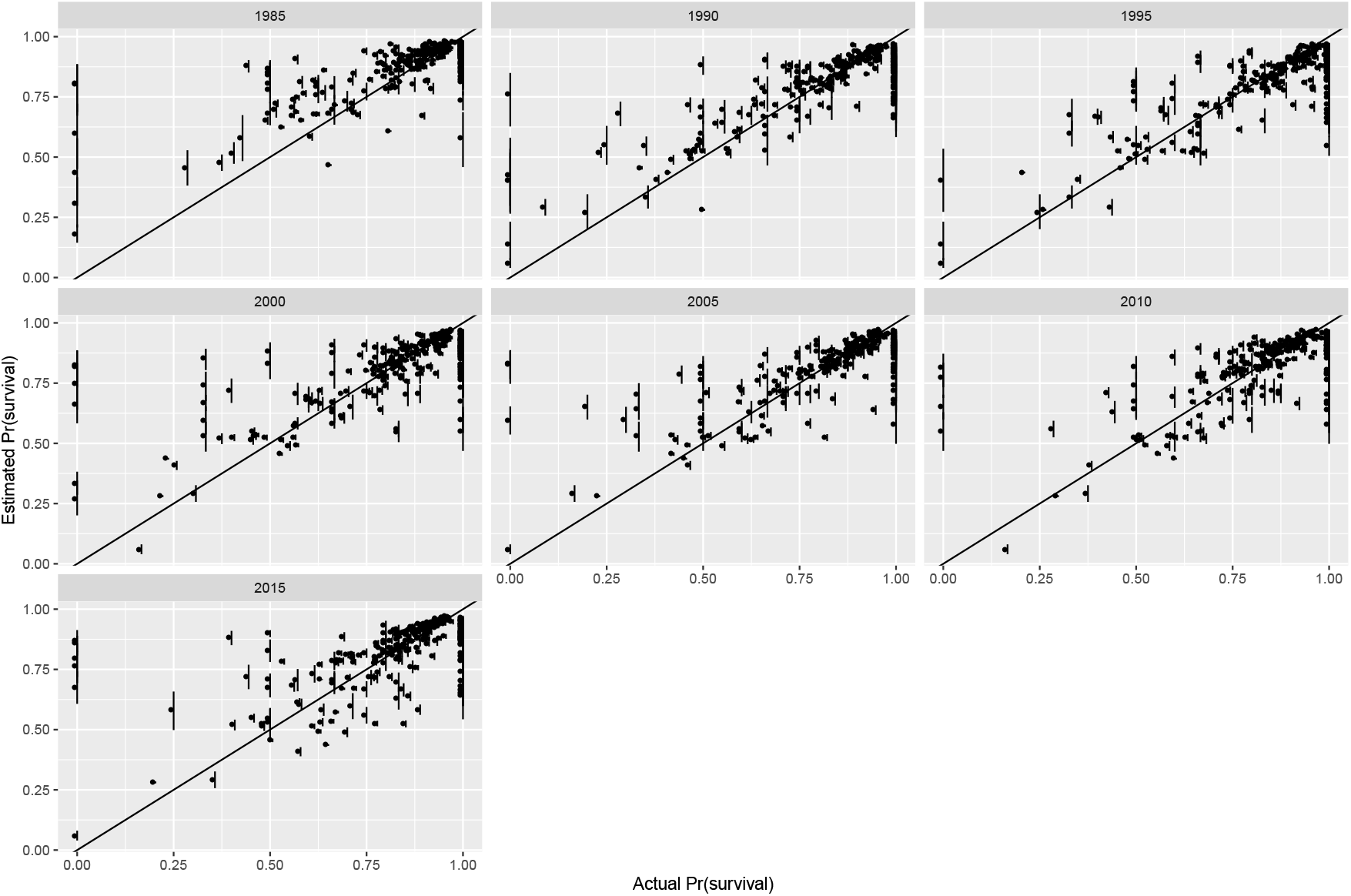
Posterior predictive distribution for the birth-death model: annual survival probabilities by census year. Each panel indicates actual versus predicted annual survival probabilities for a given census year. The vertical point-range whiskers indicate 25-75% posterior quantiles; the vertical point positions indicate posterior medians.

### 0.11 Estimating selection coefficients

To obtain selection coefficients, we first calculate the absolute fitness, *W*, of the *i*th species:

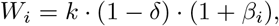

where *k* is a constant and *δ* is the estimated immigration rate. Next we calculate the relative fitness, *w_i_* of each species as:

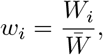

where 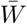 is the median absolute fitness of all species. The selection coefficient of the *i*th species is then:

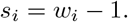

### 0.12 Testing the “demographic chance” hypothesis

We considered the possibility that the observed variance in fitness among species was due to chance differences in their demographic composition rather than their intrinsic adaptedness, but further analysis shows this not the case. To see this suppose that some species just happen to have mostly large (or old) and highly fecund individuals where others have mostly small (or young) and unproductive ones; since individual trees live for many years and can contribute offspring to multiple censuses, the former species would appear to have higher fitness than the latter. To test this “demographic chance” hypothesis we divided the BCI plot into four quadrants and re-estimated the selection parameters, *β_i_*, in each. If these quadrants are essentially independent of each other due to dispersal limitation of seeds — if they can be considered, in effect, replicates — and nevertheless show similar frequency dynamics then we can conclude that they are not governed by random demographic variation but the general properties of species. Figure S6 shows that considering all species, there is a positive, statistically significant, correlation among the *β_i_*s estimated in all four quadrants (0.48 ≥ *r* ≤ 0.67), highly so if we consider just those 57 species that are under selection in the plot as a whole (0.84 ≥ *r* ≤ 0.94). Thus, species that change substantially in frequency in one quadrant generally do so in all, which strongly suggests that they do so because of some intrinsic species-specific property rather than the chance allocation of individuals with especially high or low fecundity.

**Figure S5:**
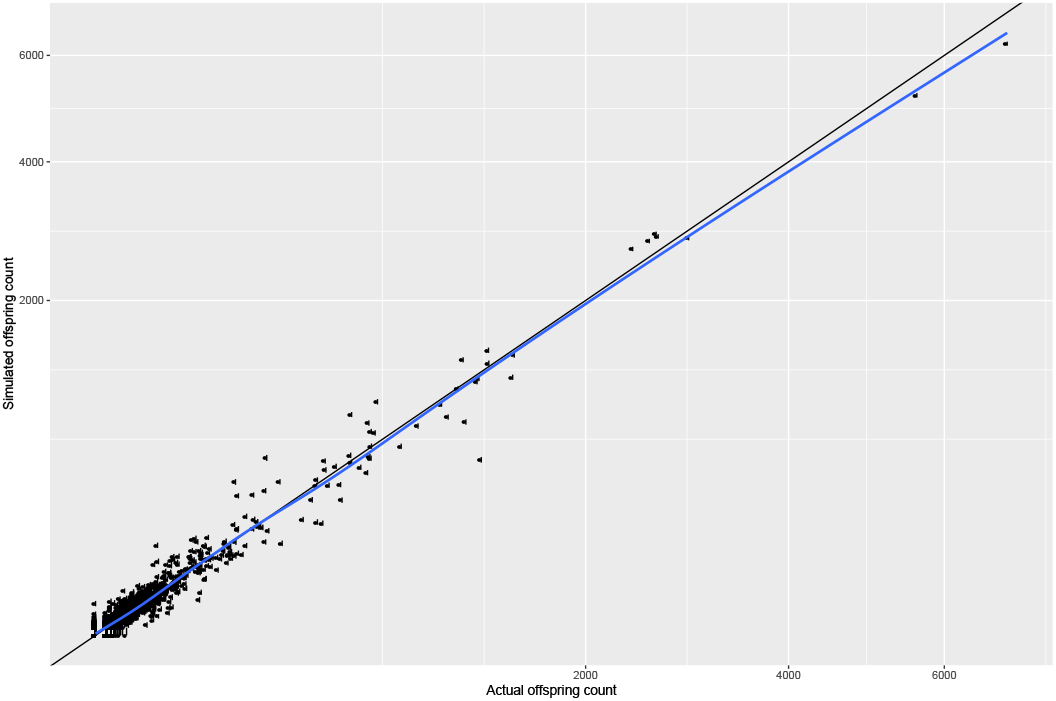
Posterior predictive distribution for the birth-death model: recruitment. The position of the vertical points indicates the posterior median number of offspring; the 25%-75% quantiles are shown but are not visually discernible. The black line indicates the actual = simulated line; the blue line indicates the linear regression of simulated on actual numbers of offspring.

### 0.13 Estimating diversity

We estimated two measures of diversity, species richness, ^0^*D*, and species evenness, ^2^*D*, the latter by means of Simpson Diversity, estimated using the `diversity` function in `R’s vegan` package converted into Hill numbers to give effective species numbers [46].

**Figure S6:**
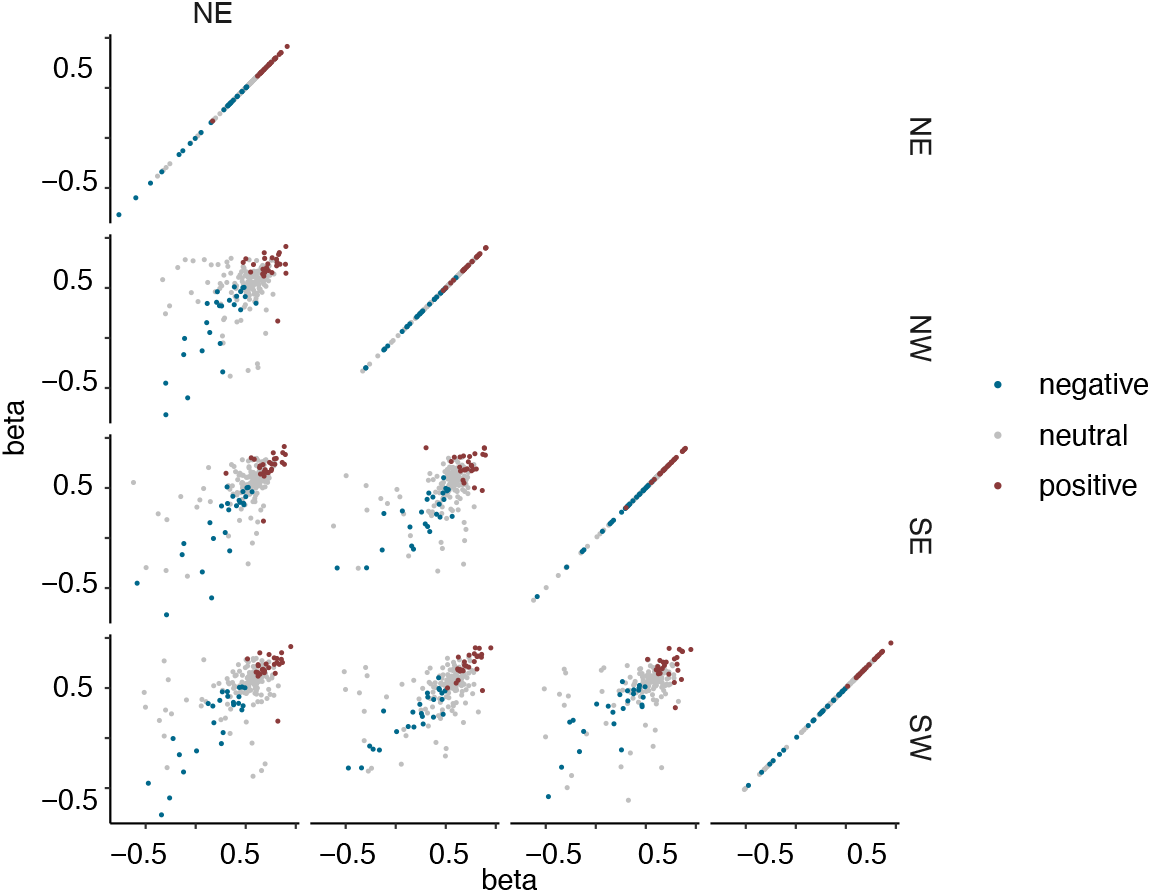
Estimates of *β_i_* in four quarters of BCI. The estimates are statistically significantly correlated when considering all species, highly so when considering only those species shown by the global analysis to be under selection (see text). This implies that the frequency dynamics that underlie the selection coefficients are indeed due to some intrinsic species-specific property rather than the chance allocation of highly fecund individuals.

**Figure S7:**
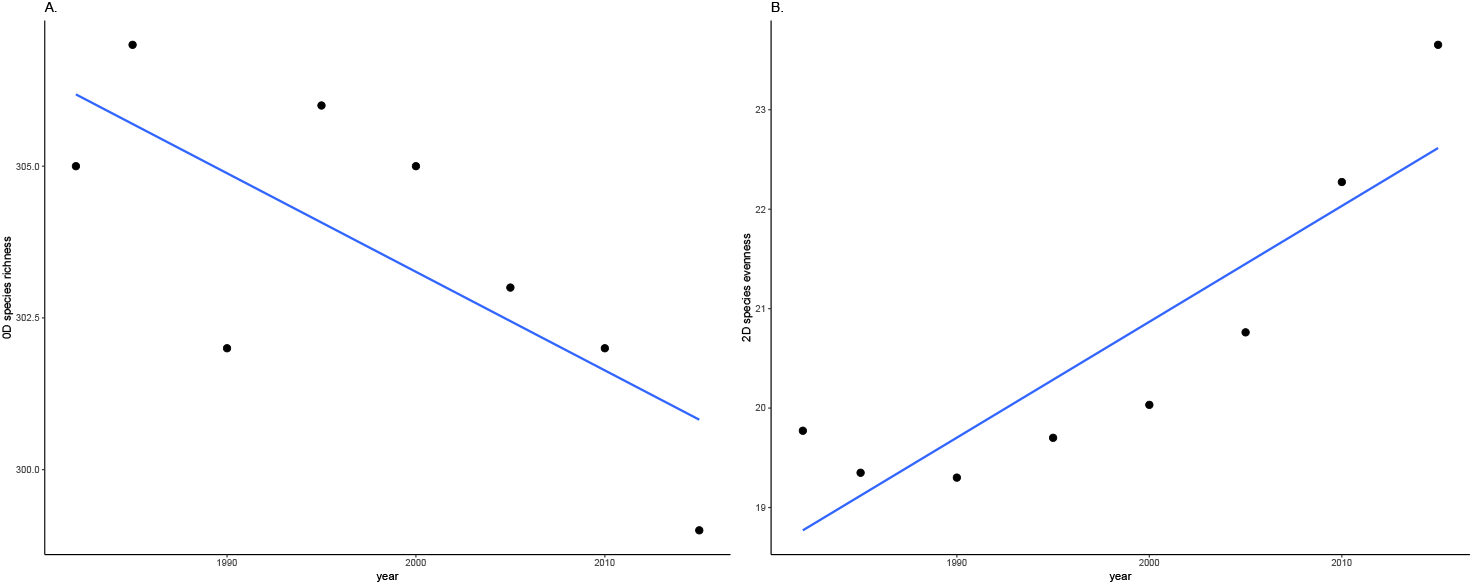
BCI diversity 1982-2015. This is based on the full dataset of 327 species recorded alive at least once. A linear model showed that species richness ^0^*D* declined significantly (*P* = 0.037) between 1982 and 2015 at a rate of −0.16 ± 0.15 species per year which implies a loss of one species every 6 years. Species evenness, ^2^*D*, increased significantly (*P* = 0.003) at a rate of 0.12 ± 0.06 effective species per year which implies a gain of one effective species every 8-9 years.

**Figure S8:**
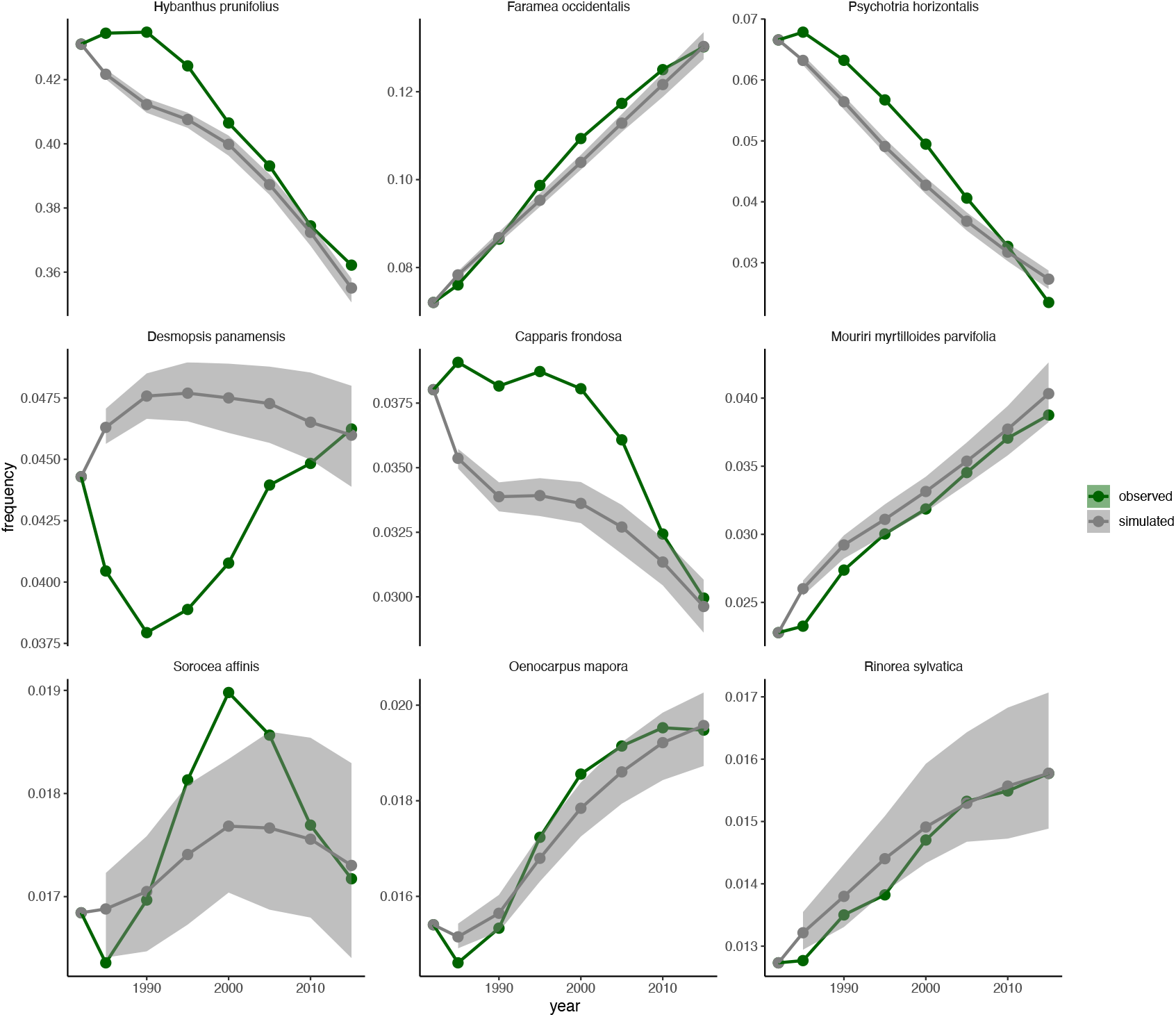
Frequency dynamics of the nine most common species: observed and simulated. The uncertainty ribbons indicate the 2.5%-97.5% simulation quantiles.

**Figure S9:**
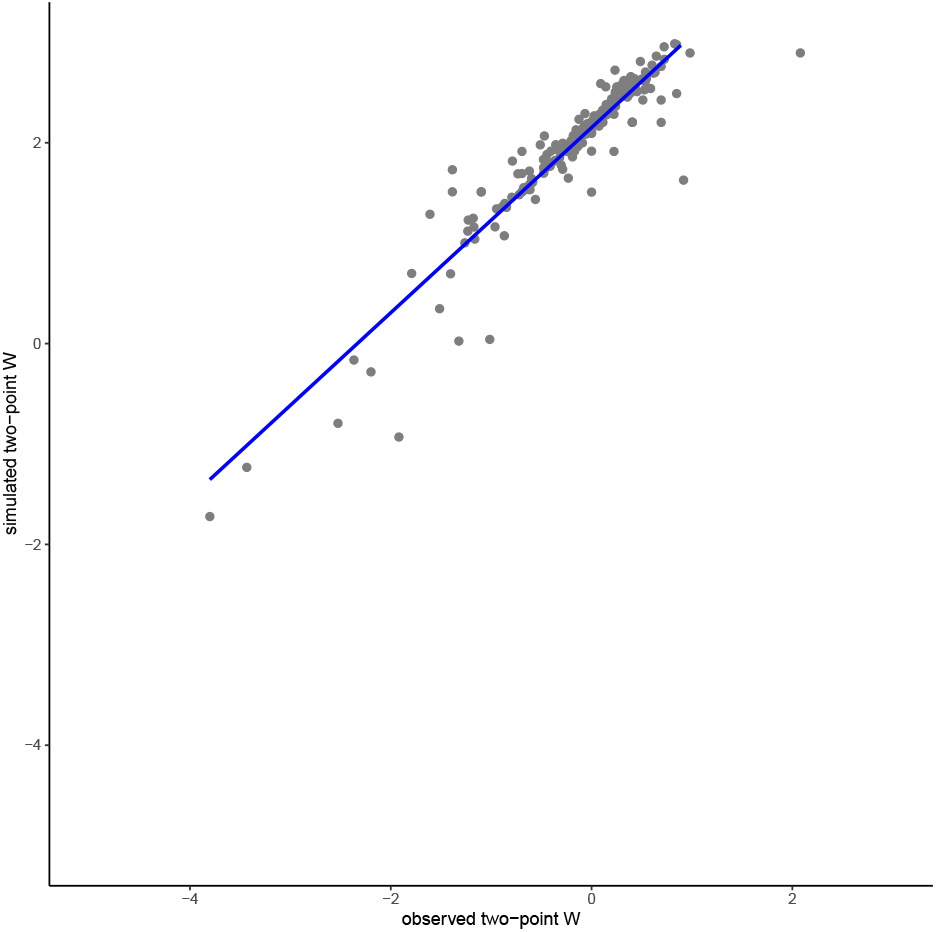
The simulations recapitulate the observed frequency dynamics of BCI species 1982–2015. We estimate the change in relative abundance of each species in the observed and simulated communities as two-point fitnesses, 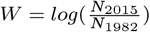. The slope of the regression is 0.922 ± 0.044 (estimate and 95%CI) as estimated by linear model, with a *r*^2^ = 0.88. We cannot estimate fitness for species not present in 1982 and 2015 so compare 228 species in all.

**Figure S10:**
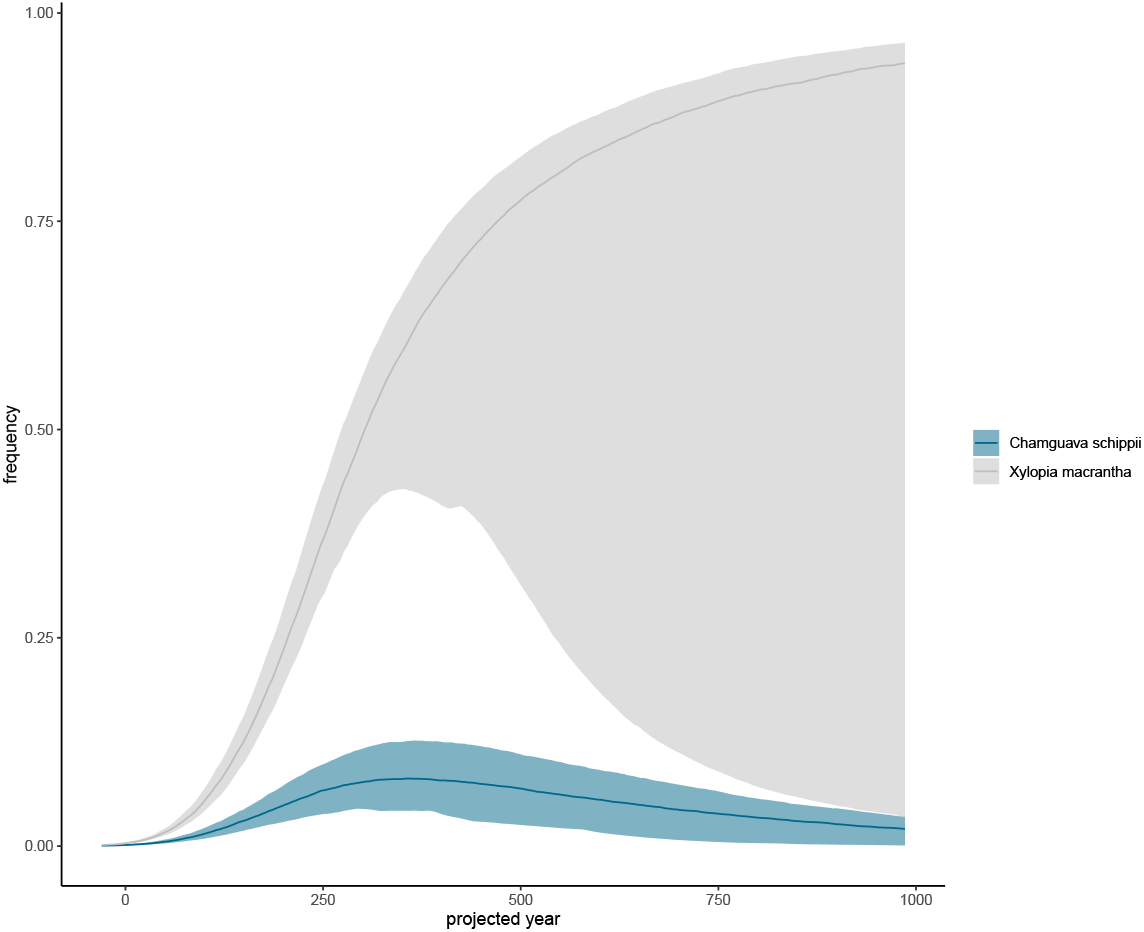
The long-term decline in species evenness in detail. We identified the two most abundant species for each of a hundred simulated communities in 483 years into the future, when the median species evenness is 3.3. These were *Chamguava schippi* and *Xylopia macrantha*. At that time they have a combined frequency of 0.83. The uncertainty intervals show the 2.5%-97.5% simulation quantiles.

## Notes

https://github.com/ben18785/bci_temporal_dynamics

### Competing Interest Statement

The authors have declared no competing interest.

